# Organization of digestive functions in the chordate gut

**DOI:** 10.64898/2026.11.24.768686

**Authors:** Arun R. Chavan, Parastou Yaghoubi, Olivia C. Langner, Fabian S. Menges, Kimberley H. Gibson, Kejue Jia, Jacob M. Musser, Scott J. Miller, Larry J. Dishaw, Ruslan Medzhitov

## Abstract

Despite their widely different anatomies, the digestive tracts of all animals must perform a set of core functions that allow the animal to acquire and utilize nutrients from the environment. Whether there are general principles that govern how these functions are spatially organized, however, remains unclear. Here we show that the digestive tract of the tunicate *Ciona robusta* is organized into pigmented and unpigmented domains specialized for nutrient uptake and metabolism, respectively. We identify the major pigment as halocynthiaxanthin, a diet-derived carotenoid that accumulates in the vacuolated, microvilli-rich epithelium of the pigmented domains, marking the absorptive regions along the gut. We find a similar spatial separation of uptake and metabolism in a distantly related tunicate, *Styela clava,* despite a markedly different pigmentation pattern, as well as in the mouse small intestine along both the anterior–posterior and crypt–villus axes. These findings suggest that the spatial separation of uptake from metabolism is a conserved principle of chordate gut organization. We propose that the underlying trade-off between these two functions may have served as an organizing force in the evolution of specialized digestive cell types and organs.

## 2. Introduction

Animal digestive systems are remarkably diverse. Most bilaterians have through-guts, cnidarians have sacs with a single opening (Steinmetz, 2019), sponges filter-feed using a unidirectional water current (Godefroy et al., 2019), and placozoans digest food externally along their ventral surface (Smith & Mayorova, 2019). Even within narrow clades, digestive tracts vary dramatically to accommodate ecological niches and feeding strategies, such as between herbivorous and carnivorous mammals (Karasov et al., 2011; Morton, 1979).

Despite their anatomical diversity, all digestive systems must perform a set of core functions that allow animals to acquire and use nutrients from their environment (Yonge, 1937). Food is first captured and broken down extracellularly by a combination of mechanical, chemical, and enzymatic processing. Exocrine cells produce the enzymes that break down specific macromolecules, such as carbohydrates, proteins, and lipids. The resulting products are taken up by absorptive cells through apical transporters. In most animals, larger particles can also be internalized by phagocytosis and digested intracellularly (Hartenstein & Martínez, 2019). Absorbed nutrients are then metabolized to derive energy, used as building blocks for biosynthesis, or stored for later use.

In certain lineages, these functions have been allocated to specialized structures during evolution. In vertebrates, for example, extracellular digestion, absorption, and metabolism are broadly allocated to the pancreas, the intestine, and the liver, respectively. Other animal lineages lack these organs entirely, and in at least some of them comparable functions are partitioned among regions or cell types along the gut (Annunziata et al., 2019). Echinoderm and tunicate digestive tracts, for instance, consist of a bulbous glandular stomach followed by an intestine, whereas cephalochordates and many protostomes have tubular digestive tracts with one or more midgut diverticula (Morton, 1979). Moreover, the midgut and hindgut, where digestion and absorption primarily take place, are organized less consistently across bilaterians than the sharply demarcated foregut (Vianello et al., 2026). The same set of functions is thus organized differently in the digestive tracts across animal lineages, but whether there are general principles that govern how these functions can, and cannot, be spatially arranged in the digestive tract is unknown.

We noticed that the digestive tract of adult *Ciona robusta* (formerly *Ciona intestinalis* type A; hereafter *Ciona*) displays a visually striking pattern of pigmentation, with sharply defined pigmented and unpigmented regions occurring at stereotyped positions across animals of different sizes. Although this pigmentation pattern is visible in previous anatomical publications (Iguchi et al., 2023), it has not been formally described or characterized. We therefore asked whether the pigmentation pattern reflects an underlying functional organization of the digestive tract and whether this organization could reveal evolutionarily conserved principles governing how digestive functions are spatially arranged.

Here, we show that the *Ciona* digestive tract is functionally organized into domains specialized for nutrient uptake or metabolism, and that this organization maps onto pigmented and unpigmented regions, respectively. This functional separation is not unique to *Ciona*: in a distantly related tunicate *Styela clava* (hereafter *Styela*), despite a markedly different overall pattern of digestive tract pigmentation, pigmented and unpigmented regions similarly distinguish uptake and metabolism programs. We further find that these functional modules remain spatially separated in *Mus musculus* (hereafter mouse) despite substantial differences in digestive tract organization, extending across spatial scales from the local crypt–villus axis to the organ-wide anterior–posterior axis of the intestine. Together, these findings suggest that the spatial separation of uptake and metabolism represents a deeply conserved principle of digestive tract organization, maintained across chordates through major changes in anatomy.

## 3. Results

### 3.1. The Ciona robusta digestive tract exhibits spatially distinct domains

We observed a striking pattern of pigmentation along the anterior–posterior axis of the *Ciona* digestive tract. The stomach (S) was orange, followed by alternating unpigmented and pigmented intestinal (I) segments, designated I1–I4 from anterior to posterior: unpigmented I1, pigmented I2, unpigmented I3, and pigmented I4. These regions were separated by sharp and reproducible boundaries, suggesting spatial compartmentalization of the digestive tract. Similar to the villi in the mammalian digestive system, the stomach epithelium exhibited major and minor folds. Pigmentation was spatially restricted to the tips of these folds, whereas the crypt-like regions remained unpigmented. This pattern is indicative of an additional axis of organization within the stomach (Fig. 1a).

**Fig. 1.**
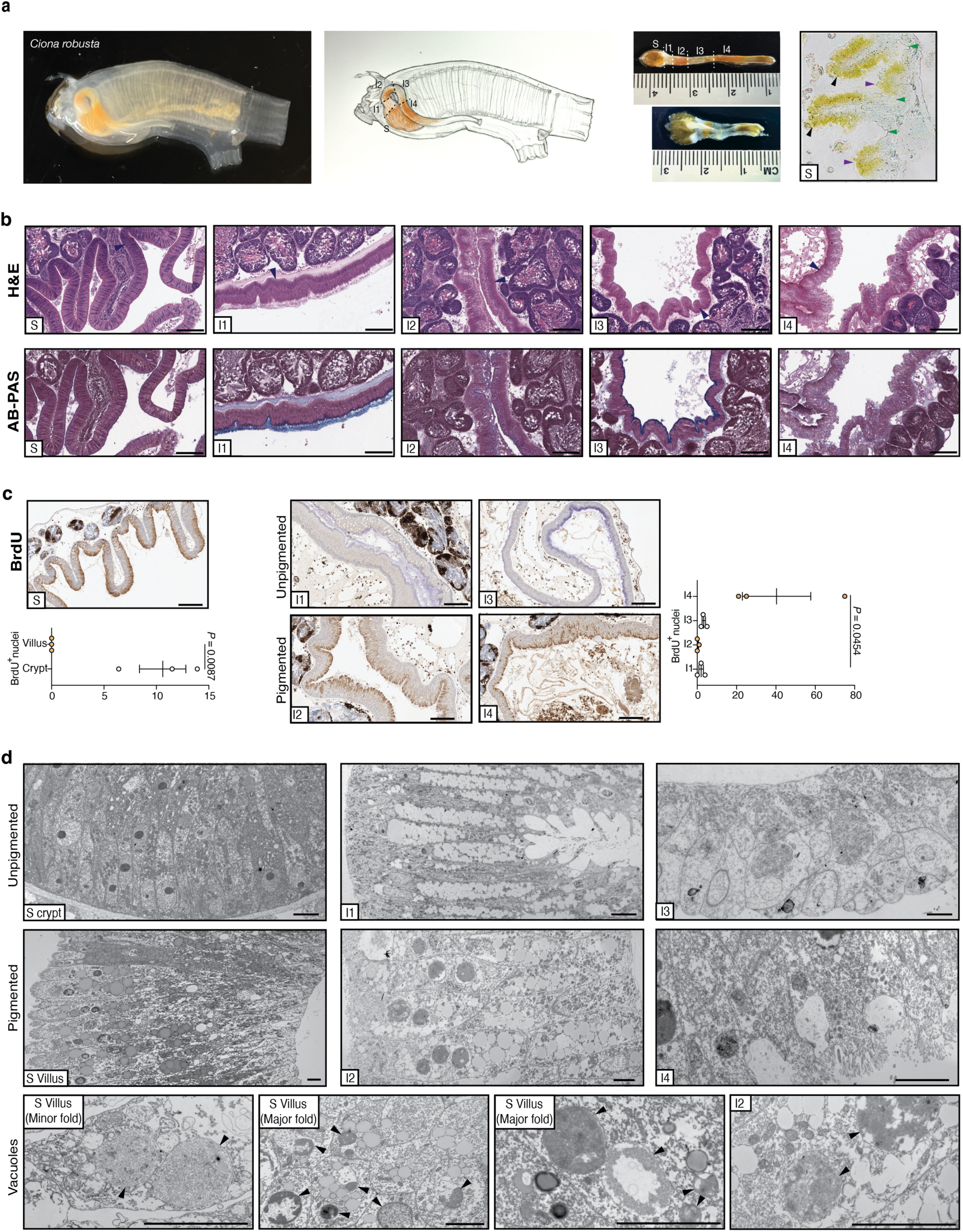
Pigmentation marks distinct functional domains along the *Ciona* digestive tract. **a,** Digestive system of *Ciona*, with the stomach (S) and four intestinal segments (I1–I4) highlighted. I1 and I3 are unpigmented, whereas I2 and I4 are pigmented. Stomach crypts and minor and major folds are indicated by green, purple and black arrowheads, respectively, in an unstained fresh-frozen OCT section. **b,** Representative H&E and AB-PAS staining of stomach and intestinal sections. Arrowheads denote the epithelial layer. **c,** BrdU incorporation in the stomach and intestinal segments following feeding of animals (*n* = 3) with 0.3 mg ml⁻¹ BrdU in artificial seawater for 24 h. **d,** Electron micrographs of stomach crypts and folds and intestinal segments. Vacuoles enriched in pigmented regions are indicated by black arrowheads. For **c,** data are mean ± s.e.m., dots represent biological replicates (≥4 measurements/replicate); *P* values by unpaired two-tailed *t*-tests. Scale bars, 100µm **(b, c)** and 5µm **(d)**.

Histological analysis by hematoxylin and eosin (H&E) staining further distinguished these regions. Pigmented regions generally appeared clearer and less eosinophilic than adjacent unpigmented regions. The stomach epithelium also contained interspersed, morphologically distinct cells with dense basophilic cytoplasm, which may represent a secretory population. Alcian blue–periodic acid–Schiff (AB–PAS) staining revealed a blue-stained mucus layer covering the epithelium in the unpigmented intestinal segments (I1 and I3) (Fig. 1b). These observations suggest functional specialization between pigmented and unpigmented regions. Proliferation was also spatially restricted: proliferative cells in the stomach were predominantly localized to the crypt-like regions, whereas intestinal proliferation was highest in the distal I4 segment (Fig. 1c).

Electron microscopy (EM) further revealed distinct subcellular organization of the pigmented domains. Pigmented regions (tips of the stomach folds, I2, and I4) contained abundant microvilli and prominent vacuolar structures (Fig. 1d), with the extensive vacuolation consistent with the clearer appearance of these regions by H&E staining (Fig. 1b). These vacuolar structures were distinct from the previously described (Thomas, 1970b) mucin-containing vacuoles observed in the unpigmented regions (I1 and I3) (Supplementary Fig. 1) which correspond to the AB–PAS-positive regions. Together, these findings reveal that the *Ciona* digestive tract has a defined spatial organization, in which pigmentation marks domains with distinct histological, secretory, proliferative, and subcellular properties.

### 3.2. Diet-derived carotenoids mark vacuolated domains of the digestive tract in Ciona

Because functional specialization appeared to align with the sharply delineated pigmentation pattern, we next sought to determine the identity and localization of the digestive tract pigment. Liquid chromatography-mass spectrometry (LC-MS) analysis identified the predominant pigment as halocynthiaxanthin (Fig. 2a), a carotenoid previously identified in the body wall of another tunicate, *Halocynthia* (Matsuno et al., 1984; Matsuno & Ookubo, 1981). This identification was further supported by the observed and previously published (Yamano et al., 1995) proton nuclear magnetic resonance (¹H NMR) spectra (Supplementary Fig. 2) and by high-resolution mass spectrometry (HRMS-ESI+), with an observed [M+H]⁺ of *m/z* 599.4080, closely matching the calculated value of 599.4095 for halocynthiaxanthin (C₄₀H₅₅O₄⁺). Three additional minor peaks were also detected. Although their identities could not be confirmed by NMR owing to their low abundance and instability, their masses and retention time were consistent with related carotenoid metabolites (Fig. 2b). LC–MS analysis across the digestive tract further showed that all four pigment-associated peaks, including halocynthiaxanthin and the three minor components, were consistently present across all pigmented segments (Fig. 2c).

**Fig. 2.**
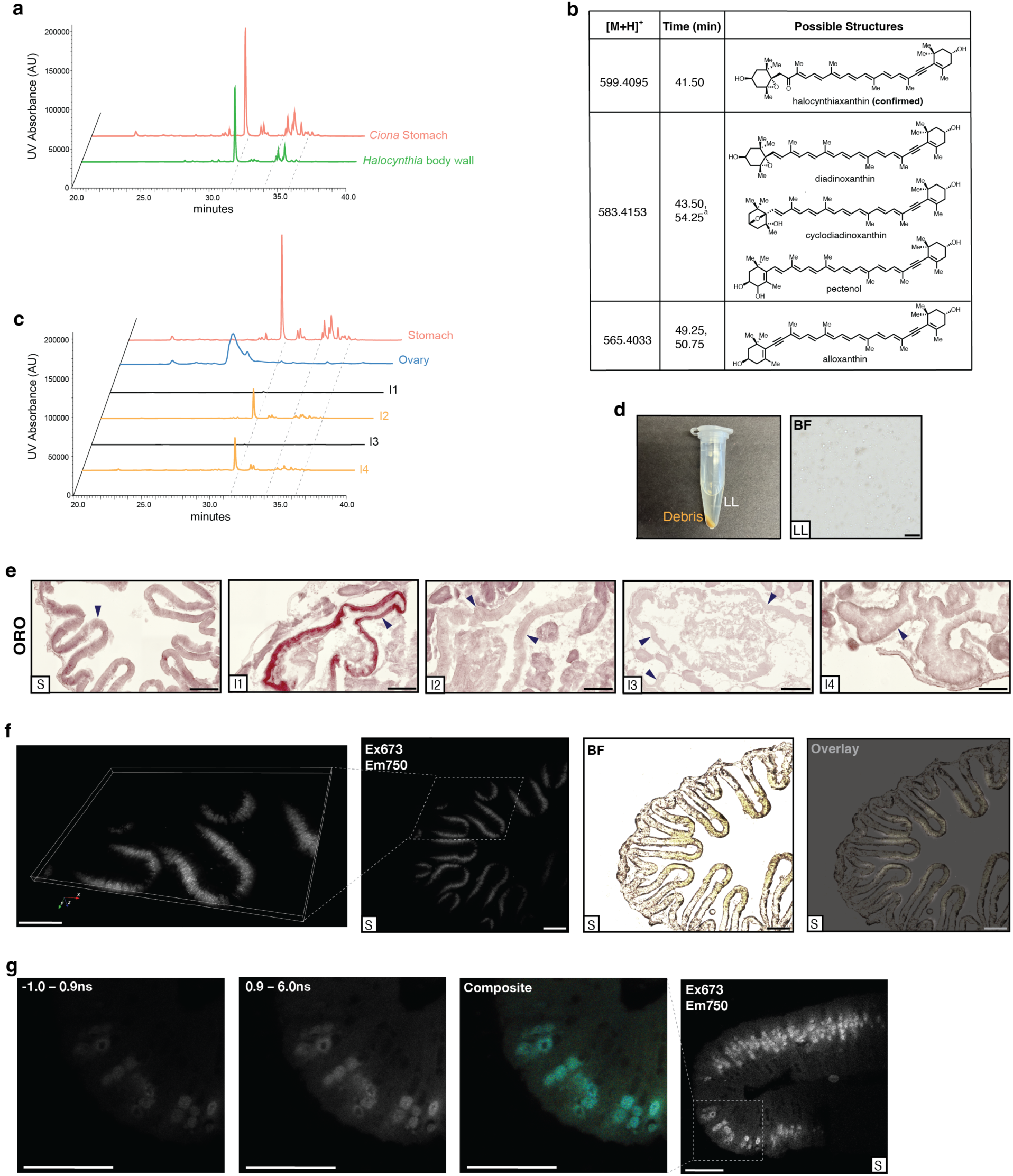
Characterization of *Ciona* digestive tract pigmentation identifies carotenoid-like molecules localized to vacuoles. **a,** 45-min gradient HPLC chromatograms monitored at 455 nm for crude extracts from the *Ciona* stomach (pink) and *Halocynthia* body wall (green). **b,** Masses of major pigmented bands in the crude *Ciona* stomach extract, based on the 70-min gradient HPLC. ᵃThis peak could alternatively correspond to an ion at *m/z* 550.6279 ([M+H]⁺) although we were unable to identify a known metabolite of the relevant class matching this mass. **c,** 45-min gradient HPLC chromatograms monitored at 455 nm for crude extracts from the *Ciona* stomach (pink), ovaries (blue), unpigmented intestinal regions (black) and pigmented intestinal regions (yellow). **d,** Representative bright-field (BF) images of the lipid-droplet layer (LL) and isolated lipid droplets from the *Ciona* stomach. **e,** Representative Oil Red O (ORO) staining of *Ciona* stomach and intestinal sections. Arrowheads denote the epithelial layer. **f,** Confocal image of a *Ciona* stomach section excited at 673 nm and detected in the far-red channel (emission maximum ∼750 nm), overlaid on the corresponding bright-field image. Pigments showed maximal autofluorescence at this excitation and emission. **g,** Zoomed-in view of a stomach fold showing pigment-containing cells. Images were acquired using TauContrast, which separates fluorescence signals based on photon arrival time (fluorescence lifetime). Composite image is built by photons detected in two discrete temporal gates (arriving times). Scale bars, 50µm **(d),** 100µm **(e–f)**, 25µm **(g)**.

Because the ovaries of *Ciona* are also strongly pigmented, we considered whether digestive tract pigmentation might originate through vertical transfer of maternally deposited pigment. However, the predominant ovarian pigment differed from that detected in the digestive tract (Fig. 2c). This observation argues against direct transmission from the ovary and instead suggests that digestive pigmentation is acquired from dietary sources.

We next asked whether the hydrophobic nature of carotenoids might explain their spatial distribution through association with lipid droplets. Isolated lipid droplets, however, lacked any pigmentation, and the pigment instead precipitated with the insoluble debris (Fig. 2d). Moreover, electron microscopy revealed lipid droplets throughout the intestine. The lipid droplets were not restricted to the pigmented region; on the contrary, they were most abundant in the unpigmented I1 region. Notably, I1 was also the only intestinal segment that showed prominent neutral-lipid staining by Oil Red O (ORO) (Fig. 2e). The discrepancy between the widespread lipid droplets observed in EM and the I1-specific ORO staining suggests further compartmentalization of metabolic processes that lead to segment-specific lipid composition.

The pigmentation exhibited intrinsic fluorescence, with excitation around 673 nm and emission in the far-red range, allowing us to examine its subcellular localization by confocal microscopy. Autofluorescent pigment was concentrated toward the apical side of epithelial cells, away from the nuclei (Fig. 2f), and frequently surrounded ring-like structures consistent with the prominent vacuoles observed by electron microscopy (Fig. 2g; Fig. 1d). These observations suggest that the pigment is associated with the apical vacuolar compartment rather than with cytoplasmic lipid droplets.

Together, these findings indicate that digestive tract pigmentation most likely reflects the regional accumulation of a diet-derived carotenoid rather than lipid storage or vertical pigment inheritance. Its enrichment within apical, microvilli-rich and vacuolated epithelial domains suggests that pigmentation marks regions specialized for intestinal uptake of dietary material.

### 3.3. *Ciona* pigmented and unpigmented segments are specialized for uptake and metabolic functions, respectively

The observations above (the dietary origin of the pigment and its apical localization in microvilli-rich domains) suggest that the pigmented regions are specialized for absorption. To further evaluate this possibility and for a comprehensive transcriptional comparison among them, we performed bulk RNA-sequencing of the four intestinal segments.

As a first step, to interpret the transcriptional differences in functional terms, we defined a hierarchical set of core functional modules that a digestive system is expected to carry out (Fig. 3). For each module, we manually curated a list of informative genes; we used mouse genes as the reference nomenclature and mapped them to their *Ciona* orthologs (Supplementary Table 1).

**Fig. 3.**
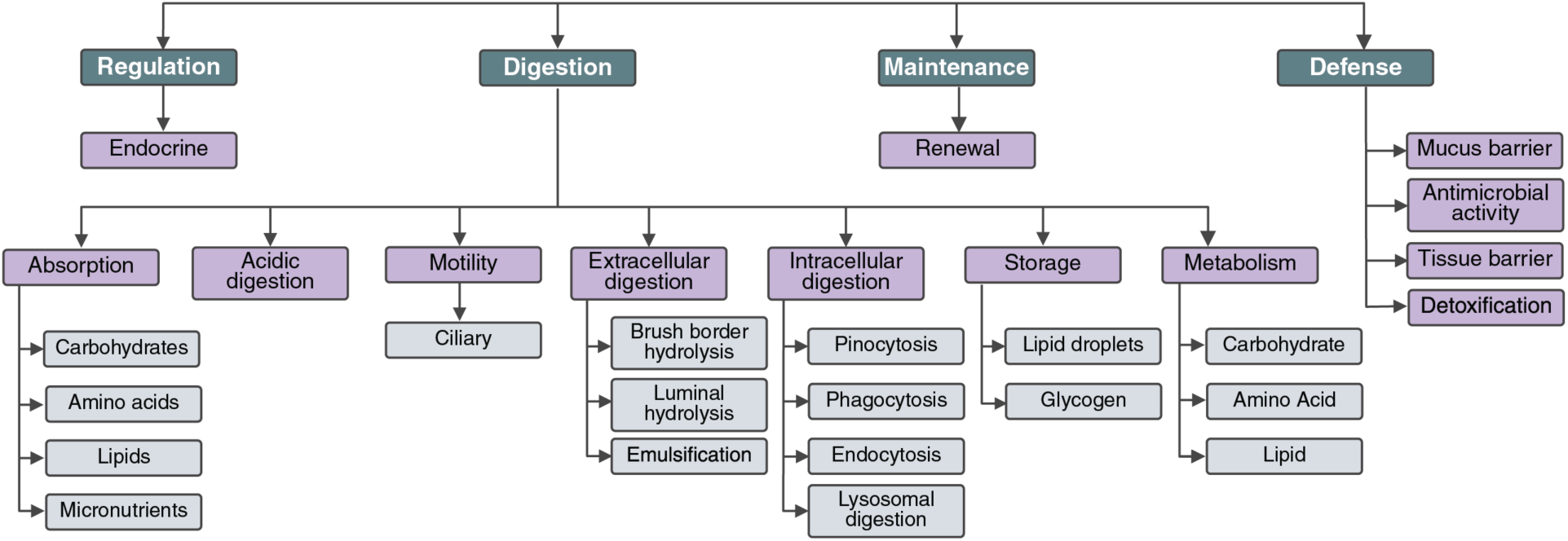
Flow chart showing the hierarchical organization of core functional modules expected to be performed by a digestive system.

Notably, the pigmented segments I2 and I4, despite not being contiguous, are transcriptionally more similar to each other than either is to its adjacent unpigmented segment (Fig. 4a). Consistent with this observation, the first principal component, explaining 41% of the variation, separates the pigmented segments from the unpigmented segments (Fig. 4b). These observations suggest underlying functional similarity between the pigmented segments.

**Fig. 4.**
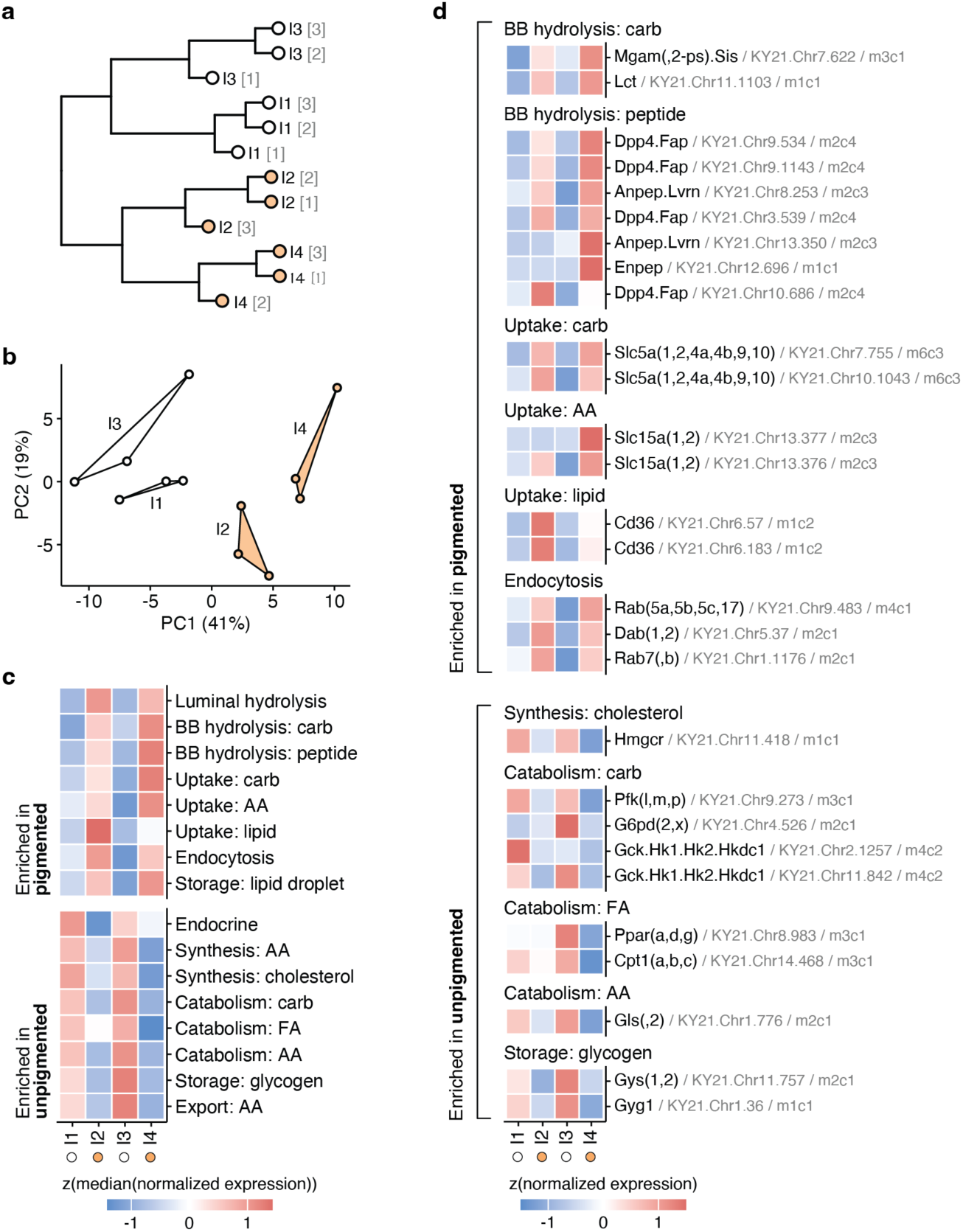
Bulk RNA-seq of *Ciona* intestinal segments shows differential deployment of uptake and metabolism functional modules. **a**, Hierarchical clustering of intestinal segments. Open and filled circles represent unpigmented and pigmented segments, respectively. **b**, PCA of intestinal segments. **c**, Expression of functional modules in the intestinal segments. The module expression is summarized as the median of L1-normalized TPM values for all genes in the module, and Z-scaled for visualization. **d**, Expression of select individual genes across intestinal segments. L1-normalized TPM values are Z-scaled for visualization. Gene labels are constructed from hierarchical orthogroups inferred with OrthoFinder (Emms et al., 2026) and follow the pattern “informative gene label / gene ID / paralog counts”. Olfactores-level hierarchical orthogroups were used. The informative gene label is derived from mouse gene symbols in the orthogroups. Symbols sharing a root gene family name are collapsed and distinguishing suffixes are listed in parentheses, e.g. Slc15a(1,2) denotes *Slc15a1* and *Slc15a2*, and symbols with distinct roots are separated by a period. Paralog counts are listed as “m[number of mouse paralogs]c[number of *Ciona* paralogs]” in the orthogroup; the mouse genes are the reference genes used to define the functional modules. Abbreviations: BB (brush border), carb (carbohydrate), AA (amino acid), FA (fatty acid).

We then quantified the expression of all functional modules in the four segments, where the expression of a functional module is the median of L1-normalized expression (i.e., each gene’s expression scaled to sum to one across the four segments) of all genes in the module. The functional modules separate clearly between segment types: modules related to nutrient uptake are enriched in the pigmented segments while those related to nutrient metabolism are enriched in the unpigmented segments (Fig. 4c). The enriched uptake-related modules include all steps in extracellular digestion from luminal hydrolysis, hydrolysis by brush border membrane enzymes, to absorption through membrane transporters. *Ciona* orthologs of apical transporter genes for all three macronutrient classes are enriched in the pigmented segments (Fig. 4d). These include the active monosaccharide transporter gene family *Slc5*; high-capacity, proton-coupled peptide importer genes *Slc15a1* (also known as PEPT1) and *Slc15a2*; and fatty acid importer *Cd36*. Enrichment of endocytosis genes implies intracellular digestion in the pigmented segments. In contrast, the unpigmented segments show enrichment of genes for catabolism of all three macronutrient classes. These include the ortholog of phosphofructokinase-1 (*Pfk*) family genes, *G6pd*, and hexokinases for carbohydrate catabolism, the ortholog of *Cpt1* family and the ortholog of *Ppar* family genes for fatty acid catabolism, and the ortholog of glutaminase genes for amino acid catabolism. The unpigmented segments are enriched, additionally, for genes involved in glycogen synthesis for storage as well as synthesis of cholesterol.

### 3.4. Segregation of uptake and metabolism is conserved across tunicates and vertebrates

We then tested if the functional segregation between uptake and metabolism is restricted to *Ciona* intestine or conserved more broadly across chordates.

First, we examined the gut of another tunicate *Styela clava*. These two genera, *Ciona* (Phlebobranchia) and *Styela* (Stolidobranchia), bracket most of tunicate diversity, sharing a common ancestor with all tunicates except appendicularians (Fig. 5a). Anatomically, the digestive tract of *Styela* has a similar structure to that of *Ciona* with an anterior bulbous stomach followed by an intestine. The pigmentation pattern of the stomach is also similar with pigmented villi and unpigmented crypts. The pigmentation pattern of the intestine, however, is different. Unlike *Ciona*, where pigmented and unpigmented segments alternate, *Styela* pigmentation grades from bright anteriorly (I2–I4) to pale posteriorly (I5–I6), with an unpigmented first segment (I1) (Fig. 5b). Accordingly, the first PC (47% of variation) aligns with the anterior–posterior axis; the second PC (21%) aligns with pigmentation (Fig. 5c).

**Fig. 5.**
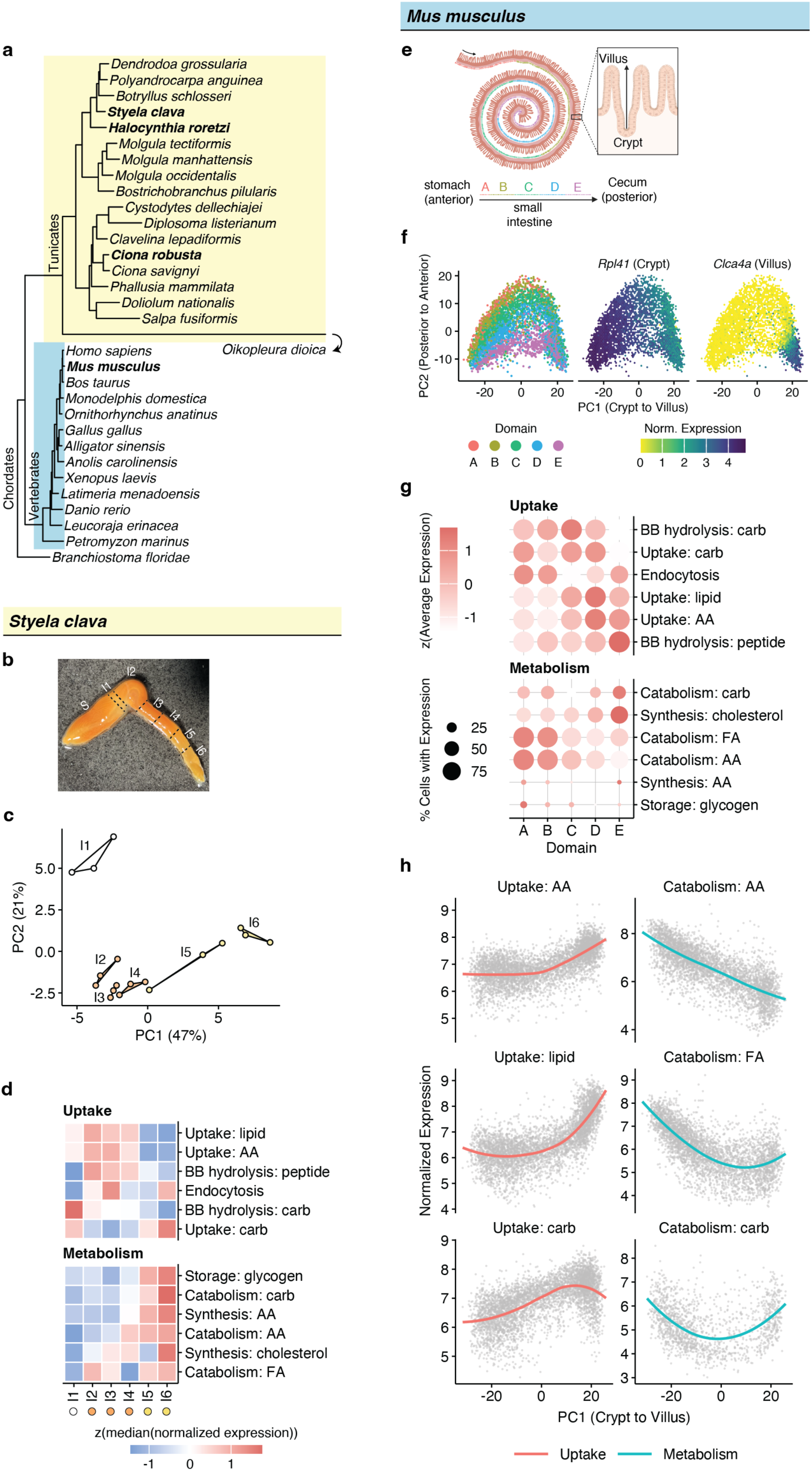
Separation of uptake and metabolism programs is conserved across tunicates and vertebrates. **a**, Phylogeny of chordates with names of species used in this study emphasized in bold. The tree was obtained from Delsuc et al. (2018) and was replotted after pruning to the subset of species shown. The tree did not include *Styela clava*, therefore the tip for its congener *Styela plicata* was labelled as *Styela clava*. Panels **b–d** correspond to bulk RNA-seq of *Styela*. **b**, Digestive tract of *Styela* with the stomach (S) and six intestinal segments (I1–I6) highlighted. The segment I1 is unpigmented, I2–I4 have bright pigmentation, I5 and I6 have pale pigmentation. **c**, PCA of bulk RNA-seq data from *Styela* intestinal segments. The fill color of the circles represents pigmentation. **d**, Expression of functional modules in intestinal segments. Only uptake and metabolism functional modules are shown. As in Fig. 4c, the expression of a functional module is calculated as the median of L1-normalized TPM values for all genes in the module, and Z-scaled for visualization. Panels **e–h** correspond to reanalysis of mouse enterocyte single-cell RNA-seq data from Zwick et al. (2024). **e**, An illustration of the mouse small intestine, which varies along two axes: anterior-posterior and crypt-villus. The anterior-posterior length is divided into 5 domains A–E defined by Zwick et al. (2024). **f**, PCA of mouse enterocytes. Anterior–posterior domains A–E are distinguished by color in the left panel. Expression of marker genes for enterocytes from the crypt (*Rpl41*) and villus (*Clca4a*) is plotted in the PC space in the right panels. **g**, Expression of functional modules along the anterior-posterior domains. The expression of a functional module in a cell is calculated by aggregating counts for all genes in the module per cell followed by normalization. **h**, Expression of functional modules for uptake and catabolism of macronutrients along the crypt–villus axis denoted by the first principal component. Abbreviations: BB (brush border), carb (carbohydrate), AA (amino acid), FA (fatty acid).

Despite the different organization of the pigmented regions, the functional organization is consistent with *Ciona*. As in *Ciona*, functional modules for nutrient uptake are enriched in the brightly pigmented segments (I2–I4), while metabolism modules are enriched in the pale segments (I5–I6) (Fig. 5d). The carbohydrate uptake module is an exception to this clean separation: when summarized across all genes, it appears to be enriched in the unpigmented segments. However, this reflects the behavior of lowly expressed paralogs and passive transporters (*Slc2* family). For active monosaccharide transporters (*Slc5a* family), the most highly expressed paralogs peak in the pigmented segments, consistent with the overall pattern of enriched uptake in pigmented segments, while lowly expressed paralogs show the reverse pattern (Supplementary Fig. 3).

Next, we asked whether this pattern extends beyond tunicates to vertebrates. Using single-cell RNA-seq data from Zwick et al. (2024), we assessed the functional specialization of enterocytes along the two axes of mouse intestine organization: anterior–posterior and crypt–villus (Fig. 5e). Principal components analysis of enterocytes shows that crypt–villus and anterior–posterior dimensions are the primary and secondary axes of variation corresponding to PC1 and PC2, respectively (Fig. 5f). Accordingly, we use PC1 as a continuous proxy variable for assessing functional specialization along the crypt–villus axis. Along the anterior–posterior axis, nutrient uptake modules are enriched in segments C and D, whereas metabolism modules are enriched in segments A, B, and E (Fig. 5g). This functional division is also seen along the crypt–villus axis. The uptake modules are enriched toward the villi whereas the catabolism modules are enriched toward the crypt, with a gradual change along the axis (Fig. 5h). Along both axes, anterior–posterior and crypt–villus, the functional separation is clearer for lipids and amino acids than for carbohydrates. As in *Styela*, this reflects the behavior of passive monosaccharide transporter (*Slc2a5*), which is broadly expressed; the active transporter (*Slc5a1*) follows the pattern seen for other macronutrients with enrichment in segments C and D (Supplementary Fig. 3).

## 4. Discussion

Our histological and transcriptional analyses reveal that the *Ciona* digestive tract is spatially organized into functionally distinct domains, broadly separating regions specialized for nutrient absorption from those enriched in metabolic programs. A similar organization is observed in the digestive tract of *Styela* and mouse, suggesting that spatial separation of nutrient acquisition and metabolic processing may represent a conserved principle of digestive tract organization. This functional segregation between uptake and metabolism likely reflects an underlying trade-off.

In both *Ciona* and *Styela* yellow pigmentation provides a striking marker of absorptive domains. These regions are transcriptionally enriched for fatty acid, amino acid, and active carbohydrate transporters, as well as alkaline phosphatases, previously described to be associated with intestinal nutrient absorption (Burighel & Milanesi, 1973). The identification of halocynthiaxanthin as the predominant *Ciona* pigment, a compound most likely derived from the dietary fucoxanthin found in algae (Maoka, 2021), further supports the idea that pigmentation marks regions with high uptake activity. Whether these carotenoids contribute to organismal physiology in tunicates remains unclear. Their accumulation may simply reflect dietary uptake followed by intracellular sequestration, or they may have functional roles, such as signaling or structural functions, within epithelial cells. This possibility is particularly interesting given that carotenoid metabolism in vertebrates generates signaling molecules such as retinoic acid (Von Lintig et al., 2021). Additionally, studies in model lipid membranes show that these hydrophobic molecules can associate with vesicular and plasma membranes and influence their biophysical properties (Gruszecki & Strzałka, 2005). Thus, although pigmentation provides a useful readout of absorptive organization in tunicates, whether these dietary carotenoids are merely retained products of absorption or have biological function remains an open question.

Another prominent feature of the pigmented regions is the abundance of apical vacuoles. Similar vacuolated epithelial cells have been described in other tunicates, including *Botryllus schlosseri* (Burighel & Milanesi, 1973), *Ciona intestinalis* (Thomas, 1970a), and Pyuridae (Degail & Lévi, 1964), and have been associated with absorptive functions (Burighel & Cloney, 1997). Notably, however, in these tunicates, as well as in *Ciona*, the vacuoles are structurally heterogeneous. Some contain membranous or myelin-like structures, whereas others contain small vesicles or amorphous electron-dense material, suggesting that they may represent functionally distinct compartments. Vesicular structures are also present in unpigmented regions, where some appear to be mucin-rich, further suggesting functional diversity among these compartments. The formation and functions of these diverse vacuoles remain poorly understood and will require further biochemical characterizations.

Functional regionalization within the intestine has been investigated largely through the lens of nutrient absorption. In mouse, enterocytes are zonated along both the anterior–posterior and crypt–villus axes, specializing in absorption of different nutrient molecules (Moor et al., 2018; Zwick et al., 2024). In the juvenile *Ciona* intestine, Iguchi et al. (2023) found a repetitive and zonal expression of absorptive genes, with some regions (compartments c6 and c8, corresponding to our I1 and I3) lacking absorptive gene expression. However, their analysis could not reveal what these regions do. We curated a hierarchical set of gene modules necessary for a functioning digestive system, covering nutrient uptake, metabolism, storage, regulation, maintenance, and defense functions. These modules allowed us to survey functional specialization beyond absorption and across a complete set of functional programs. Analyzing the transcriptomic data through these modules showed that I1 and I3 are enriched in metabolic programs, revealing a previously unrecognized trade-off between nutrient uptake and metabolism.

The recurrence of this functional separation across tunicates and vertebrates, despite differences in digestive tract organization, suggests that it arises from intrinsic physiological constraints rather than lineage-specific adaptations. Spatially separating nutrient uptake from metabolic utilization may confer several advantages. First, it allows nutrient-absorbing cells to deliver nutrients faithfully to other tissues rather than consuming them locally. For example, mammalian enterocytes preferentially fuel themselves with glutamine, glutamate, and aspartate (Alpers, 2000; Windmueller & Spaeth, 1980), allowing much of the absorbed dietary glucose to reach the portal vein for subsequent metabolic processing by the liver. Second, this separation may allow nutrient uptake to respond to environmental availability without being constrained by, or interfering with, intracellular metabolic and biosynthetic programs. Metabolic intermediates can regulate nutrient transport through feedback mechanisms; for instance, in hepatocytes which are tasked with metabolic processing, glucose-6-phosphate promotes GLUT1 internalization (Sasson et al., 1997) and degradation (Qualls-Histed et al., 2023), limiting uptake. However, glucokinase, a key enzyme for hepatic glucose utilization is not subject to this inhibition (Ashcroft et al., 2023).

Spatial segregation of functions is not limited to incompatible digestive functions. Barrier defense and tissue renewal are also spatially segregated in the mouse small intestine along both the anterior–posterior and crypt–villus axes. Goblet cells, the major producers of mucus, increase in abundance from anterior to posterior, reaching their highest abundance in the ileum (Ermund et al., 2013; Singh, 1971), whereas, proliferation capacity decreases from anterior to posterior (Holt et al., 1983; Parker et al., 2017). Similarly, the mature mucus-secreting goblet cell subpopulation is enriched in the villus compartment (Dolan et al., 2022; Gustafsson et al., 2021), while the proliferative intestinal stem cells are confined to the crypt compartment (Cheng & Leblond, 1974).

One evolutionary pathway to resolution of functional trade-offs is the gradual specialization of cells toward one function at the expense of the other (Shoval et al., 2012). This results in a continuum of variation, as seen along the crypt–villus axis in enterocytes and the porto–central axis in hepatocytes, bounded by functional archetypes, i.e., cells maximally specialized for one function (Adler et al., 2019, 2023; Korem et al., 2015). The same functional trade-off, and its resolution, can operate at different levels of organization. The archetypes within one cell type can crystallize during evolution into distinct cell types. For instance, enterocytes and hepatocytes can be conceptualized as sister cell types that arose through individuation of the uptake and metabolism archetypes of an ancestral multifunctional digestive cell type. Similarly, the diversification of digestive organs in vertebrates represents specialization along the same functional fault lines: intestine and liver specialized for uptake and metabolism, respectively. We propose that this functional trade-off, operating across scales and taxa, is a major organizing force in chordate digestive system evolution.

## 5. Methods

### 5.1. Animals

Adult *Ciona robusta*, formerly *Ciona intestinalis* type A (Pennati et al., 2015), were acquired from either M-REP (San Marcos, CA, USA) or Marinus Scientific (Lakewood, CA, USA) and kept at room temperature in artificial seawater (ASW) prepared with Instant Ocean Sea Salt. The animals were used for tissue collection within one day. Adult *Styela clava* were collected by the Marine Resources Center at Marine Biological Laboratory (Woods Hole, MA, USA). Animals were dissected on site and tissues were flash-frozen or fixed as appropriate for further processing.

### 5.2. Histology

Dissected animals were fixed overnight in 4% paraformaldehyde (PFA) in ASW at 4 °C. For H&E and AB-PAS staining, samples were dehydrated in 70% ethanol and embedded in paraffin. Staining was performed by either the Yale Pathology Core or HistoWiz. For Oil Red O (ORO) staining, samples were cryoprotected in 30% sucrose, embedded in Tissue-Tek optimal cutting temperature (OCT) compound (Sakura) and sectioned to 10 µm thickness using a cryostat. ORO staining was performed as previously described (https://mousepheno.ucsd.edu/pdfs/1_Oil_Red_O_HC.pdf) using Oil Red O solution (Sigma-Aldrich, O1391).

### 5.3. *In Vivo* BrdU Proliferation Assay

Animals were fed 0.3 mg ml⁻¹ BrdU (Cayman Chemical, 15580) in ASW and sacrificed 24 h later. Fixed samples were embedded in paraffin as described above, and sections were stained for BrdU by HistoWiz (Abcam, ab6326).

### 5.4. Transmission Electron Microscopy

Tissue preparation and fixation were adapted from previously published protocols (Ballarin et al., 2010; Dishaw et al., 2011). Briefly, *Ciona* stomach and intestinal tissues were dissected into rings, opened longitudinally and cut into approximately 1 × 1 mm pieces. Tissues were immediately transferred to ice-cold filtered seawater containing 1.5% glutaraldehyde and 0.2% osmium tetroxide and fixed for 1 h on ice with gentle intermittent agitation every 1–5 min, followed by washes in filtered seawater, 0.2 M sodium cacodylate (pH 8.0) and water. Tissues were stained *en bloc* with 2% uranyl acetate for 1 h in the dark and dehydrated through a graded ethanol series (50%, 70% and 90% ethanol, followed by three changes in anhydrous 100% ethanol). Tissues were transitioned through 1:1 (v/v) propylene oxide:100% ethanol, followed by two changes of pure propylene oxide and then 1:1 (v/v) propylene oxide : EPON overnight. Tissues were subsequently infiltrated with EPON overnight, embedded in fresh EPON and polymerized at 60 °C for 48 h.

Semithin (250 nm thick) sections were cut on a Leica UC7 Ultramicrotome, stained with Toluidine Blue via light microscopy to ascertain quality and orientation of the intestine and gut. Afterwards, ultrathin sections (55 nm thick) were cut and mounted on formvar-coated Copper slot grids, followed by post-staining with 2% uranyl acetate and Reynolds’ lead citrate (Reynolds, 1963).

Samples were imaged on an 80 kV Tecnai G2 Spirit BioTWIN Transmission electron microscope (Thermofisher Scientific, Hillsboro, OR, USA). Images were acquired using a NanoSprint15 MKII camera (AMT Imaging, Woburn, MA) using the AMT Capture Engine, Software version 7.0.2.5.

### 5.5. Lipid droplet isolation

Lipid droplets were isolated using a commercially available kit (Abcam, ab242290) according to the manufacturer’s instructions. Briefly, stomachs pooled from multiple animals were dissected and minced with scissors. Samples were incubated with 400 µl “Reagent A” on ice for 10 min, followed by the addition of 1600 µl 1× “Reagent B” and incubation on ice for an additional 10 min. Samples were then homogenized using a Dounce homogenizer with five strokes each using pestles A and B. The homogenate was centrifuged at 20000 × *g* for 3 h. The lipid droplet fraction was recovered from the supernatant, while cellular debris remained in the pellet. Isolated lipid droplets were examined by light microscope.

### 5.6. Crude pigment extraction and characterization by LC-MS

To obtain crude pigment extracts, the stomach or individual intestinal segments were dissected and gently rubbed against 3mM cellulose chromatography paper (Whatman, 3030-347) to release and absorb the pigment. The paper was then immersed in 100% methanol for several minutes to recover the pigments in methanol. Extracts from more than 10 animals were pooled for each sample. Samples were then centrifuged at maximum speed for 5 min to remove insoluble debris.

Mass spectrometric measurements were performed on crude extracts using a Shimadzu Scientific Instruments QToF 9030 LC–MS system equipped with a Nexera LC-40D XS UHPLC, consisting of a CBM-40 Lite system controller, DGU-405 degasser unit, two LC-40D XS UHPLC pumps, SIL-40C XS autosampler and CTO-40S column oven. UV data were collected using a Shimadzu Nexera HPLC/UHPLC Photodiode Array Detector SPD-M40 over a range of 190–800 nm. Mass spectra were subsequently recorded using the quadrupole time-of-flight (QToF) 9030 mass spectrometer. Samples were maintained at 4 °C in the autosampler compartment.

For small-molecule analysis, 5–10 µl of each solution was injected into a sample loop and separated on a Shim-pack Scepter C18-120 column (1.9 µm, 2.1 × 100 mm) equilibrated at 40 °C in the column oven. A binary gradient was used with HPLC-grade CHROMASOLV water containing 0.1% formic acid as solvent A and HPLC-grade CHROMASOLV acetonitrile containing 0.1% formic acid as solvent B.

For the 45-min gradient, the flow rate was maintained at 0.400 ml min⁻¹. The eluent was held at 95% A and 5% B from 0–2 min, followed by a linear gradient to 2% A and 98% B from 2–37 min. The eluent was then held at 2% A and 98% B from 37–42 min, rapidly returned to 95% A and 5% B from 42–42.01 min, and maintained at this composition until 45 min.

For the 70-min gradient, the flow rate was maintained at 0.400 ml min⁻¹. The eluent was held at 95% A and 5% B from 0–2 min, followed by a linear gradient to 50% A and 50% B from 2–20 min and then to 2% A and 98% B from 20–60 min. The eluent was held at 2% A and 98% B from 60–67 min, rapidly returned to 95% A and 5% B from 67–67.01 min, and maintained at this composition until 70 min.

The ionization source was run in DUIS mode, with the electrospray needle held at +4.5 kV. Nebulizer gas was maintained at 2 l min⁻¹, heating gas flow at 10 l min⁻¹ and the interface at 300 °C. Dry gas was maintained at 10 l min⁻¹, the desolvation line at 250 °C and the heating block at 400 °C.

Mass spectra were recorded over a range of 100–2000 *m/z* in positive-ion mode. MS/MS fragmentation data were acquired using data-dependent acquisition (DDA) for all precursors between 200 and 1000 *m/z*, with a collision energy of 40 CE (±20 spread), and fragments were recorded between 50 and 1000 *m/z*.

Measurements and data post-processing were performed using LabSolutions 5.135 Realtime Analysis and PostRun.

### 5.7. Major pigment isolation and characterization by ^1^H NMR

To isolate the major pigmented compound for further characterization, the crude *Ciona* stomach extract was concentrated *in vacuo* and purified by normal-phase flash column chromatography using 60 Å SiO_2_ gel (32–62 µm), with mobile phase composition and gradient (1–20% methanol/dichloromethane and 100% hexanes).

The purified pigment was characterized by nuclear magnetic resonance spectroscopy (NMR). ^1^H NMR spectra were recorded on an Agilent 800 MHz spectrometer at ambient temperature. NMR solvents were purchased from Cambridge Isotope Laboratories and used without further purification. Deuterated solvents were stored at ambient temperature and were used immediately after opening. Spectra were processed using MestReNova 14.2.0 using the automatic phasing and polynomial baseline correction capabilities. Additional manual phasing was done as necessary. Splitting was determined using the automatic multiplet analysis function with manual intervention as necessary.

### 5.8. Autofluorescence confocal imaging

OCT sections (10 µm) were prepared as described above, mounted in 90:10 glycerol:water and imaged using a Leica STELLARIS 8 confocal microscope (Leica Microsystems) equipped with TauSense. Maximum excitation and emission for pigment autofluorescence was determined by scanning across the entire range of the 405 nm diode laser and the White Light Laser (WLL) (excitation range: 405 nm and 440–790 nm; detection range: 410–850 nm). TauContrast was used to improve image quality. Images were analyzed using Aivia 16 (Leica Microsystems) software.

### 5.9. Genomic data

For *Ciona*, the HT19 (Satou et al., 2019) genome assembly with the KY21 (Satou et al., 2022) gene models, transcriptome, and proteome sequences were downloaded from the Ghost database (http://ghost.zool.kyoto-u.ac.jp/download_ht.html). The mitochondrial genome, which is not included in HT19, and its gene annotations were retrieved from Ensembl release (v113) of the KH reference genome (Satou et al., 2008). For *Styela*, the RefSeq assembly GCF_013122585.1 (Wei et al., 2020) and its annotations were downloaded from NCBI.

Functional annotation of protein coding genes in both species was improved or supplemented by annotating them with eggNOG-mapper (Cantalapiedra et al., 2021) and PROST (Kilinc et al., 2023). In addition, for *Styela* functional annotation from KEGG database was also used. Gene orthology was inferred with OrthoFinder (Emms et al., 2026). Digestive functional module genes (curated as mouse genes) for *Ciona* and *Styela* were identified by orthology to mouse genes, with manual curation using their functional annotation (above) for genes lacking an ortholog call.

### 5.10. Bulk RNA-sequencing

#### 5.10.1. Sample collection

For *Styela*, the digestive tract was dissected out, segmented, collected in STAT-60 reagent (AMSBIO, CS-502), and flash-frozen. In enterogonid tunicates the intestine is surrounded by a network of testicular tubules (Kocot et al., 2018; Millar, 1953). Therefore, for *Ciona*, the digestive tract was dissected out and separated from the surrounding testis tissue before collecting it in STAT-60 reagent for flash-freezing. The intestine was cut open longitudinally and the layer of gut epithelium was peeled from the testis tissue with fine forceps under a dissecting microscope.

#### 5.10.2. RNA extraction

Samples were homogenized in STAT-60 reagent (AMSBIO, CS-502) using a Bead Ruptor Elite homogenizer (Omni International), followed by chloroform extraction. Total RNA was purified using the RNeasy Mini Kit (Qiagen, 74106) with on-column DNase I digestion. RNA concentration was quantified using a NanoDrop 8000 spectrophotometer.

#### 5.10.3. Library preparation and sequencing

For *Ciona* samples, library preparation and sequencing were performed by the Yale Center for Genome Analysis (YCGA). For *Styela* samples, libraries were prepared and indexed using the NEBNext Ultra II Directional RNA Library Prep Kit (NEB, E7760L) according to the manufacturer’s instructions. Library DNA concentrations were measured using the Qubit High Sensitivity DNA Assay Kit and Qubit Fluorometer (Life Technologies, Q32854). Pooled libraries were paired-end sequenced on an Illumina NextSeq 2000 using P2 100-cycle v3 reagents (Illumina, 20046811).

#### 5.10.4. Analysis

Sequencing reads were pseudo-aligned to the respective transcriptome of each species and expression was quantified as transcripts per million (TPM) (Wagner et al., 2012) using kallisto (Bray et al., 2016). All subsequent analyses were performed with R version 4.5.1 (R Core Team, 2025).

Before downstream analysis, we cleaned the datasets by removing three sets of genes. First, despite separating the *Ciona* gut epithelium from testis during dissection (see above), some contamination remained. We removed testis-specific genes from the dataset to prevent them from influencing downstream analyses. A total of 623 testis-enriched genes were identified, using *Ciona* testis single-cell RNA-seq data from Li et al. (2022), and removed. Second, due to the close proximity of the stomach and I1 segment in both *Ciona* and *Styela*, these segments cannot be cleanly separated during dissection. Consequently, some stomach-enriched genes appear as I1-enriched when only the intestine samples are considered. To identify stomach-derived transcripts in segment I1, we normalized each gene’s TPM values across all segments (stomach and I1–I4 for *Ciona*; stomach and I1–I6 for *Styela*) to unit sum (L1-normalization) and computed the median normalized expression per segment across replicates. Genes with median normalized stomach expression above 0.75, i.e., genes whose expression is overwhelmingly stomach-derived, were classified as stomach spillover and removed (306 genes in *Ciona* and 113 in *Styela*). Third, we removed marginally expressed genes that did not exceed 5 TPM in at least one intestinal segment in every replicate.

All further analyses were done with these cleaned datasets and only including the intestinal segment samples. To make gene expression patterns comparable across genes with different absolute expression levels, TPM values were L1-normalized across the intestinal segments. The expression of a functional module was summarized as the median of L1-normalized expression of all genes in that module, and it was Z-scaled for visualization. PCA was performed using the ‘prcomp()’ function with options ‘center = TRUE’ and ‘scale. = FALSE’, and hierarchical clustering was performed using the ‘pvclust()’ function (Suzuki & Shimodaira, 2006) with options ‘method.dist = correlation’, ‘method.hclust = average’, and ‘nboot = 1000’ using the L1-normalized data.

### 5.11. Reanalysis of single-cell RNA-sequencing data

Single-cell RNA-sequencing count matrices and metadata for mouse small intestinal epithelial cells from Zwick et al. (2024) were downloaded from NCBI GEO (accession GSM6077219) and analyzed using Seurat version 5.3.1 (Hao et al., 2024). The 30 intestinal segments were grouped into five anterior–posterior domains (A–E) as defined by Zwick et al. Data were log-normalized, and the 2000 most highly variable genes were identified and used for PCA after scaling. Clustering was performed using 30 principal components with default parameters. Clusters were annotated by their marker genes, and only enterocyte clusters were retained and re-clustered for further analysis. For functional module analysis, module expression was calculated by summing counts for all genes in the module per cell and log-normalizing.

## 6. Acknowledgments

We thank Susanne Gibboney for technical assistance with rearing and collecting *Ciona* juveniles, and David Ferrier for helpful discussions. This research made use of the Chemical and Biophysical Instrumentation Center (CBIC) (RRID:SCR_021738), supported by Yale University funds, and the Yale Center for Genome Analysis (YCGA), supported by the National Institute of General Medical Sciences of the National Institutes of Health under Award Number 1S10OD030363-01A1. Studies in the RM lab were supported by the Howard Hughes Medical Institute (HHMI) and the Blavatnik Family Foundation.

## 7. Data availability

All transcriptomics data have been deposited in the Gene Expression Omnibus (GEO) database under the SuperSeries accession GSE354417. *Ciona robusta* bulk RNA-seq data are available under GSE354357, and *Styela clava* bulk RNA-seq data are available under GSE354416.

## 8. Code availability

All code used for analysis is available at https://github.com/archavan/pigmentation along with code for preparation of genomic data at https://github.com/archavan/genomes.

## 9. AI usage

ChatGPT 6.2, Kimi K3, and Opus 5.5 were used for proofreading, polishing language, and clarity suggestions. All text was carefully reviewed and edited by the authors. The authors take full responsibility for all content.

---

**Supplementary Fig. 1.**
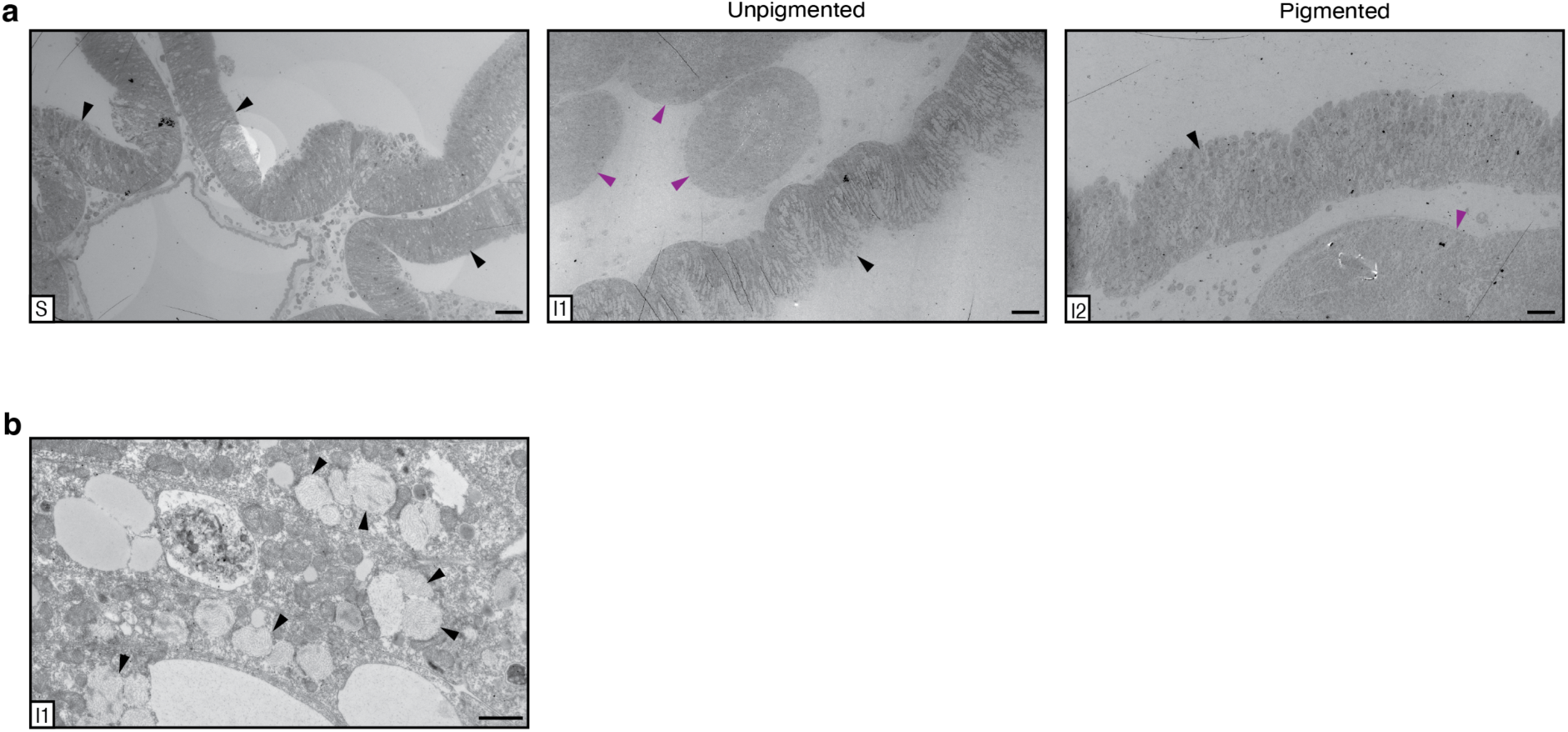
Ultrastructural organization of the *Ciona* digestive tract. **a,** Low-magnification electron micrographs of the *Ciona* stomach and unpigmented (I1) and pigmented (I2) intestinal regions. Black arrowheads indicate the intestinal epithelium, and purple arrowheads indicate the testis surrounding the digestive tract. **b,** Electron micrographs of the unpigmented I1 region, with black arrowheads indicating mucin-containing vacuoles. Scale bars, 20µm **(a)** and 1µm **(b)**.

**Supplementary Fig. 2.**
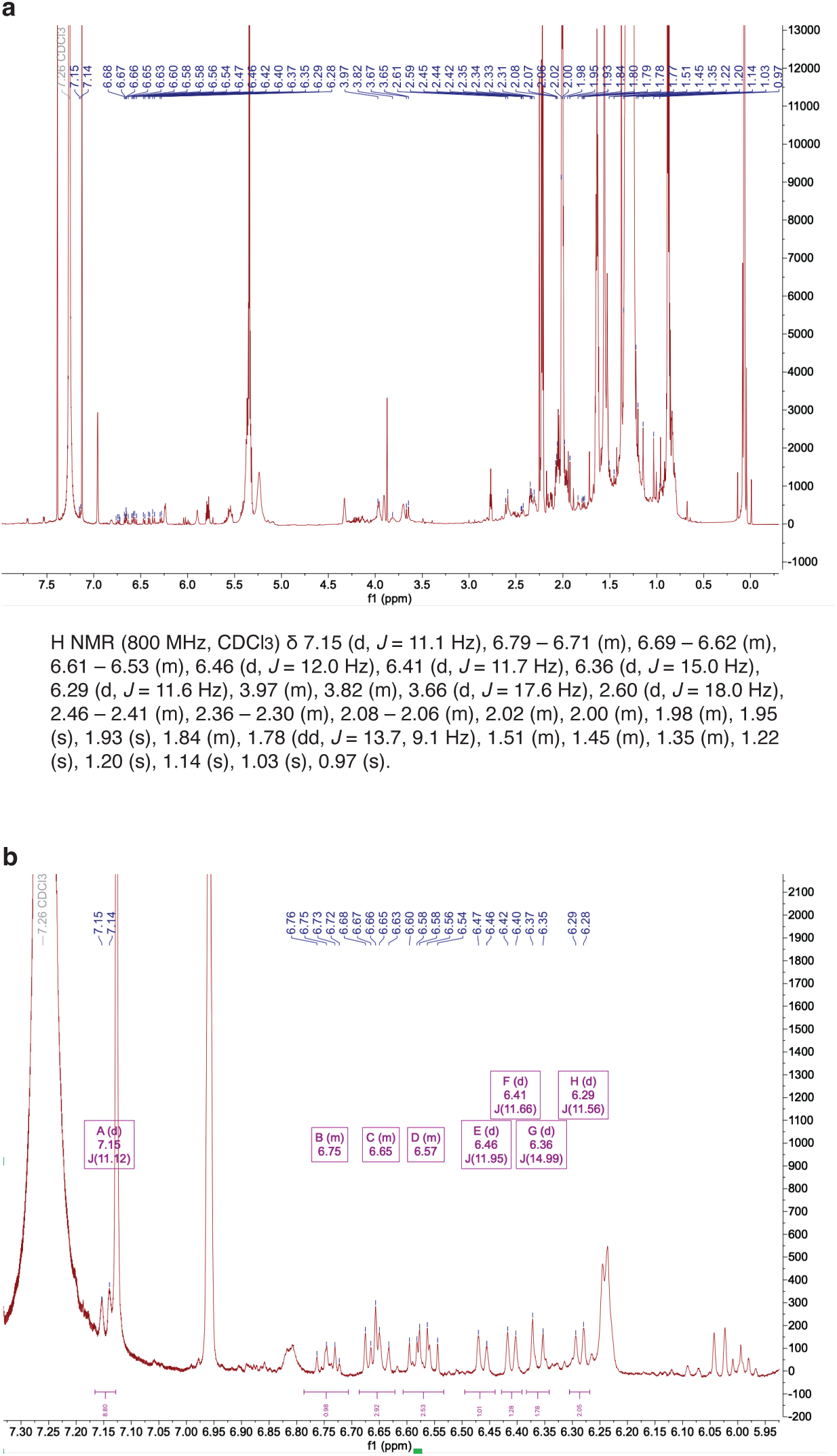
¹H NMR characterization of the major pigment isolated from the *Ciona* stomach. **a,** Full ¹H NMR spectrum of the major pigment isolated from crude *Ciona* stomach extract. Spectral data are reported as follows: chemical shift (multiplicity [singlet (s), broad singlet (brs), doublet (d), triplet (t), quartet (q), pentet (p), multiplet (m), doublet of doublets (dd), doublet of doublet of doublets (ddd), doublet of triplet of doublets (dtd), doublet of doublet of doublet of doublets (dddd), doublet of triplets (dt), triplet of doublets (td), etc.], coupling constant (*J*)). Chemical shifts (δ) are reported in ppm and *J* values in Hz. ¹H resonances were referenced to the residual solvent peak of CDCl₃ (7.26 ppm) (Fulmer et al., 2010). Integrations were inaccurate due to lack of purity and are omitted for clarity. Annotated peaks correspond to resonances reported in published spectra of purified halocynthiaxanthin (Yamano et al., 1995). **b,** Olefinic region of the ¹H NMR spectrum of the isolated pigment, showing key resonances and *J* values.

**Supplementary Fig. 3.**
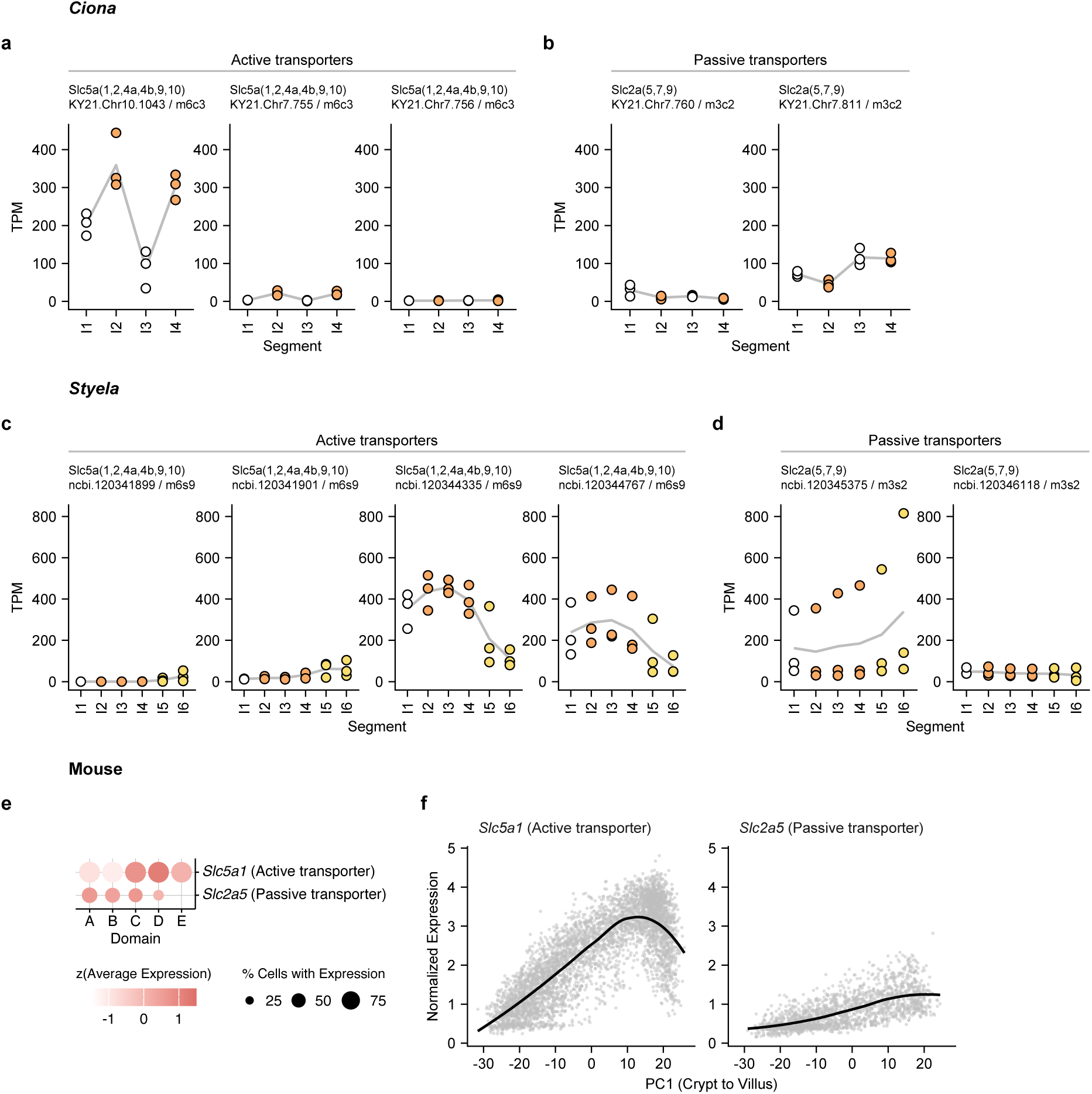
Expression of active (*Slc5* family) and passive (*Slc2* family) apical carbohydrate transporters. Expression (in TPM) of active and passive carbohydrate transporters along the intestinal segments in *Ciona* (**a** and **b**) and *Styela* (**c** and **d**). The fill color of points represents pigmentation. The grey line connects mean TPM values. The genes excluded from L1-normalization due to low expression (see Methods) are not plotted. The gene labels follow the same pattern as in Fig. 4. Briefly, they include an informative label defined as a collapsed representation of mouse gene names in the Olfactores-level orthogroup, the gene ID, and paralog count in mouse and *Ciona* or *Styela*. **e**, Expression dot plot of active and passive carbohydrate transporters along the anterior–posterior axis in mouse small intestine enterocytes; domains as defined by Zwick et al. (2024). **f**, Expression of active and passive carbohydrate transporters in mouse small intestine enterocytes along the crypt–villus axis, represented by PC1 (see Fig. 5).

**Supplementary Table 1.** Manually curated list of mouse genes for hierarchical functional modules.

| **Functional Hierarchy Level** |  |  |  | **Genes** |
|---|---|---|---|---|
| **First** | **Second** | **Third** | **Fourth** |  |
| **Regulation** | Endocrine |  |  | *Cck, Chga, Cpe, Gast, Gcg, Ghrl, Gip, Neurog3, Pcsk1, Pcsk1n, Pcsk2, Pdx1, Pyy, Scg5, Sct, Sst, Tph1* |
| **Digestion** | Acidic digestion |  |  | *Aqp4, Atp4a, Atp4b* |
|  | Motility | Ciliary |  | *Ccno, Cep41, Dnah11, Dnah5, Dnai1, Foxj1, Ift88, Lztfl1, Mcidas, Rfx3, Rsph4a* |
|  | Extracellular digestion | Emulsification | Bile acid metabolism | *Cyp27a1, Cyp7a1, Cyp7b1, Cyp8b1, Nr1h4* |
|  |  | Luminal hydrolysis |  | *Amy1, Amy2a1, Amy2a2, Amy2a3, Amy2a4, Amy2a5, Cel, Cela1, Cela2a, Clps, Cpa1, Cpa2, Ctrb1, Pgc, Pnlip, Prss1, Prss2, Ptf1a* |
|  |  | Brush border hydrolysis | Carbohydrate | *Lct, Mgam, Sis* |
|  |  |  | Peptide | *Anpep, Dpp4, Enpep, Mep1a, Mep1b* |
|  | Intracellular digestion | Pinocytosis |  | *Amn, Cubn, Lrp2* |
|  |  | Phagocytosis |  | *Actr2, Actr3, Dock1, Elmo1, Gulp1, Megf10, Mertk, Rac1* |
|  |  | Endocytosis |  | *Dab2, Msr1, Rab5a, Rab7* |
|  |  | Lysosomal digestion |  | *Acp2, Ctsb, Ctsc, Ctsd, Ctsf, Ctsh, Ctsl, Lamp1, Lamp2, Lamp3, Lamp5* |
|  | Absorption | Carbohydrate | Apical uptake | *Slc2a5, Slc5a1* |
|  |  |  | Basolateral export | *Slc2a1, Slc2a2* |
|  |  | Amino acid | Apical uptake | *Slc15a1, Slc36a1, Slc3a1, Slc6a14, Slc6a15, Slc6a19, Slc7a9* |
|  |  |  | Basolateral export | *Slc16a10, Slc3a2, Slc7a7, Slc7a8* |
|  |  | Lipid | Apical uptake | *Cd36, Npc1l1, Scarb1, Slc27a4* |
|  |  |  | Intracellular transport | *Fabp1, Fabp2, Fabp5, Fabp6, Npc1, Npc2* |
|  |  |  | Reesterification | *Acat1, Acat2, Dgat1, Dgat2, Lpcat3, Mogat1, Mogat2* |
|  |  |  | Chylomicron assembly | *Apoa1, Apoa4, Apob, Mttp, Sar1b* |
|  |  |  | Basolateral export | *Abca1* |
|  |  | Micronutrient | Uptake | *Heph, Slc11a2, Slc23a1, Slc40a1, Slc46a1* |
|  | Metabolism | Carbohydrate | Catabolism | *G6pc1, G6pd, Gck, Hk1, Hk2, Hk3, Pfkl* |
|  |  |  | Synthesis | *Pck1* |
|  |  | Amino acid | Catabolism | *Arg2, Ass1, Gls* |
|  |  |  | Urea cycle | *Arg1, Bhmt, Cps1, Otc* |
|  |  |  | Synthesis | *Gls, Glul* |
|  |  | Lipid | Synthesis | *Fasn, Scd1* |
|  |  |  | Fatty acid oxidation | *Cpt1a, Hadha, Hmgcs2, Ppara* |
|  |  |  | Cholesterol synthesis | *Hmgcr, Sqle* |
|  |  |  | Cholesterol efflux | *Abca1, Abca2, Abca3, Abca5, Abca7, Abcg1, Abcg4, Abcg5, Abcg8* |
|  | Storage | Lipid droplets |  | *Cidea, Dgat1, Dgat2, Lipe, Plin1, Plin2, Plin3, Plin4, Plin5, Pnpla2* |
|  |  | Glycogen |  | *Gyg1, Gys1, Gys2, Pygl* |
| **Defense** | Detoxification |  |  | *Abcb1a, Ahr, Ces1, Cyp1a1, Cyp1a2, Cyp2j6, Cyp3a11, Gstm1, Gstp1, Nr1i2, Nr1i3* |
|  | Tissue barrier |  |  | *Cdh1, Cldn15, Cldn2, Cldn3, Cldn4, Cldn7, Ocln, Tjp1* |
|  | Mucus barrier |  |  | *Muc2, Muc5ac, Spdef, Tff1, Tff2, Tff3* |
|  | Antimicrobial activity |  |  | *Defa5, Lyg1, Lyg2, Lyz1, Lyz2, Lyz3, Reg3g* |
| **Maintenance** | Renewal | Stem cell |  | *Lgr5, Mki67, Olfm4, Pcna, Top2a* |

